# Evidence for intraflagellar transport in butterfly spermatocyte cilia

**DOI:** 10.1101/2022.12.22.521690

**Authors:** Marco Gottardo, Maria Giovanna Riparbelli, Giuliano Callaini, Timothy L. Megraw

## Abstract

In the model organism insect *Drosophila melanogaster* short cilia assemble on spermatocytes that elaborate into 1.8 mm long flagella during spermatid differentiation. A unique feature of these cilia/flagella is their lack of dependence on intraflagellar transport (IFT) for their assembly. Here we show that in the common butterfly *Pieris brassicae* the spermatocyte cilia are exceptionally long: about 40 µm compared to less than 1 µm in Drosophila. By transmission electron microscopy we show that *P. brassicae* spermatocytes display several features not found in melanogaster, including compelling evidence of IFT structures and features of motile cilia.

**Summary statement:** This work shows ultrastructural definition of the exceptionally long cilia that persist on butterfly (*P. brassicae*) spermatocytes, including evidence of intraflagellar transport, during meiotic division.

## Introduction

Cilia are organelles that project from the cell surface with key functions in physiology and development that occur widely in eukaryotes from protists to mammals (Berbari et al., 2009; Bloodgood, 2009; Buchwalter et al., 2016; Fisch and Dupuis-Williams, 2011; Gerdes et al., 2009; Ishikawa and Marshall, 2011; Jana et al., 2016; Petriman and Lorentzen, 2020; Santos and Reiter, 2008; Singla and Reiter, 2006; Vincensini et al., 2011). They are critical organelles for signal transduction and transmit sensory input from a variety of environmental and interorgan signals. Cilia are assembled from centrioles that are retooled from their role in centrosomes into basal bodies (BB) that dock at the plasma membrane and template the assembly of the microtubule-based axoneme that is surrounded by the ciliary membrane (Dawe et al., 2007). There are varieties of cilia, including motile and non-motile cilia, and flagella, which refer to motile cilia that propel cells such as ciliates or sperm cells. The ciliary membrane, while contiguous with the plasma membrane, is compartmentalized and has a unique composition of transmembrane receptors and ion channels involved in the sensory transduction of extracellular signals (Hirokawa et al., 2006; Singla and Reiter, 2006) whose conveyance is critical for physiological and developmental processes (Berbari et al., 2009; Gerdes et al., 2009; Goetz and Anderson, 2010). Underscoring its physiological importance, dysfunction in cilia assembly and trafficking dynamics causes a spectrum of diseases collectively known as ciliopathies (Garcia-Gonzalo and Reiter, 2012; Hildebrandt et al., 2011; Valente et al., 2014).

The axoneme consists of a highly ordered array of microtubules (MTs) with a 9-fold symmetry driven by the architecture of the BB (Azimzadeh and Bornens, 2007; Goetz and Anderson, 2010; Jana et al., 2016). The makeup and organization of axonemal MTs can vary depending on ciliary functions. Cilia that are motile, such as those that generate fluid flow over tissue surfaces (Essner et al., 2002; Hirokawa et al., 2006) and those responsible for cell locomotion through flagellar propulsion (Sleigh, 1991), have dynein motor complexes (“arms”) attached to the nine doublet axonemal MTs arranged around a central pair of MTs; the “9 + 2” pattern (Fisch and Dupuis-Williams, 2011; Ishikawa, 2017; Silflow and Lefebvre, 2001). Sperm axonemes are typically the 9+2 class. The axoneme of the primary cilium, on the other hand, typically lacks the central pair (has a 9 + 0 pattern) and has no dynein arms (Takeda and Narita, 2012). Another class of motile cilia have a 9 + 0 axoneme structure with dynein arms, exemplified by the motile cilia within the node of early vertebrate embryos (Essner et al., 2002; McGrath and Brueckner, 2003; McGrath et al., 2003).

Assembly of cilia and maintenance of their length depend upon vesicular trafficking and intraflagellar transport (IFT) processes (Ishikawa and Marshall, 2011; Rosenbaum and Witman, 2002; Sung and Leroux, 2013; Taschner and Lorentzen, 2016) (Ishikawa and Marshall, 2011; Rosenbaum and Witman, 2002; Sung and Leroux, 2013; Taschner and Lorentzen, 2016). Cilium components are transported between the cell body and the cilium via bidirectional trafficking of multiprotein complexes, IFT particles, that can be visualized by TEM, or whose movements can be tracked by light microscopy (Cole et al., 1998; Qin et al., 2004; Rosenbaum and Witman, 2002; Taub and Liu, 2016). Kinesin II moves IFT particles towards the ciliary tip (anterograde transport), while retrograde transport relies on cytoplasmic dynein 2 (Hou and Witman, 2015; Ou et al., 2005; Scholey, 2008).

Spermatogenesis in many insects involves pre-spermiogenic assembly of cilia in primary spermatocytes (Klein and Wolf, 1997; LaFountain, 1976; Riparbelli et al., 2020; Wolf and Kyburg, 1989; Wolf and Traut, 1987). These cilia, or cilium-like structures, persist through both meiotic divisions (Bloodgood, 2009; Riparbelli et al., 2012). In *Drosophila*, each primary spermatocyte has two pairs of centrioles, as is typical of eukaryotic cells in G2 phase. Remarkably, all four centrioles dock at the plasma membrane during the first meiotic prophase and assemble four ciliary extensions (Fritz-Niggli and Suda, 1972; Tates, 1971) referred to as cilium-like regions (CLRs) that persist through meiosis (Avidor-Reiss and Leroux, 2015; Jana et al., 2016; Vieillard et al., 2015). These cilia are short (about 700 nm long), have a 9 + 0 axoneme devoid of dynein arms, and are therefore similar in architecture to primary cilia (Riparbelli et al., 2012) and may be composed of just the cilium transition zone (Jana et al., 2018; Vieillard et al., 2016). These insect spermatocyte cilia persist through two meiotic divisions into the spermatid stage where they further assemble into the sperm axoneme (Gottardo et al., 2013). Interestingly, the process of ciliogenesis/flagellogenesis in *Drosophila* male germ cells is independent of IFT, as mutants that block IFT complex B or the kinesin-2 motor produce functional sperm (Han et al., 2003; Sarpal et al., 2003). Butterfly spermatocytes also form cilia that persist through meiotic divisions but, in contrast to *Drosophila*, butterfly spermatocytes form very long cilia and have 9+2 axonemes (Gottardo et al., 2014; Wolf and Kyburg, 2004; Yamashiki and Kawamura, 1998).

We examined the cilia in spermatocytes in *Pieris brassicae*, also known as the cabbage butterfly or large white butterfly, a common agricultural pest native to Europe, Asia, and north Africa. While our analysis confirmed that *P. brassicae* spermatocytes have long cilia that persist in meiotic division, we also found that they contain thin filamentous structures of electron dense material at the base of the CLRs and along the ciliary membrane of the elongating CLRs that appear to be analogous to the IFT trains first observed in *Chlamydomonas* (Petriman and Lorentzen, 2020; Scholey, 2003). The findings reported here for *P. brassicae* provide new insights into a potential role for IFT in insect spermatocytes and opens new questions about differential involvement of IFTs in Drosophila spermatogenesis versus other systems.

## Results

### Features of the long cilia on butterfly spermatocytes

To investigate ciliogenesis in *P. brassicae* primary spermatocytes we performed immunofluorescence staining against acetylated α-tubulin, which preferentially labels axonemal and centriolar microtubules in a wide range of eukaryotes, including the testis of lepidopteran insects (Wolf, 1996). Acetylated tubulin was detected at two pairs of adjacent dots in young primary spermatocytes, which are likely centrioles that had not yet assembled axonemes (Fig. 1A). In later stages of meiotic prophase, the four spermatocyte cilia were extensively elongated (Fig. 1 B-D). By prometaphase (Fig. 1 E) the centriole pairs had separated, and the cilia reached their maximum length of 39.2± 4.1 µm. During the first metaphase (Fig. 1 F), each meiotic spindle pole contained one pair of cilia. Together, these data are consistent with original reports of cilia projecting from dividing spermatocytes in *P. brassicae* (Gottardo et al., 2014).

**Figure 1.**
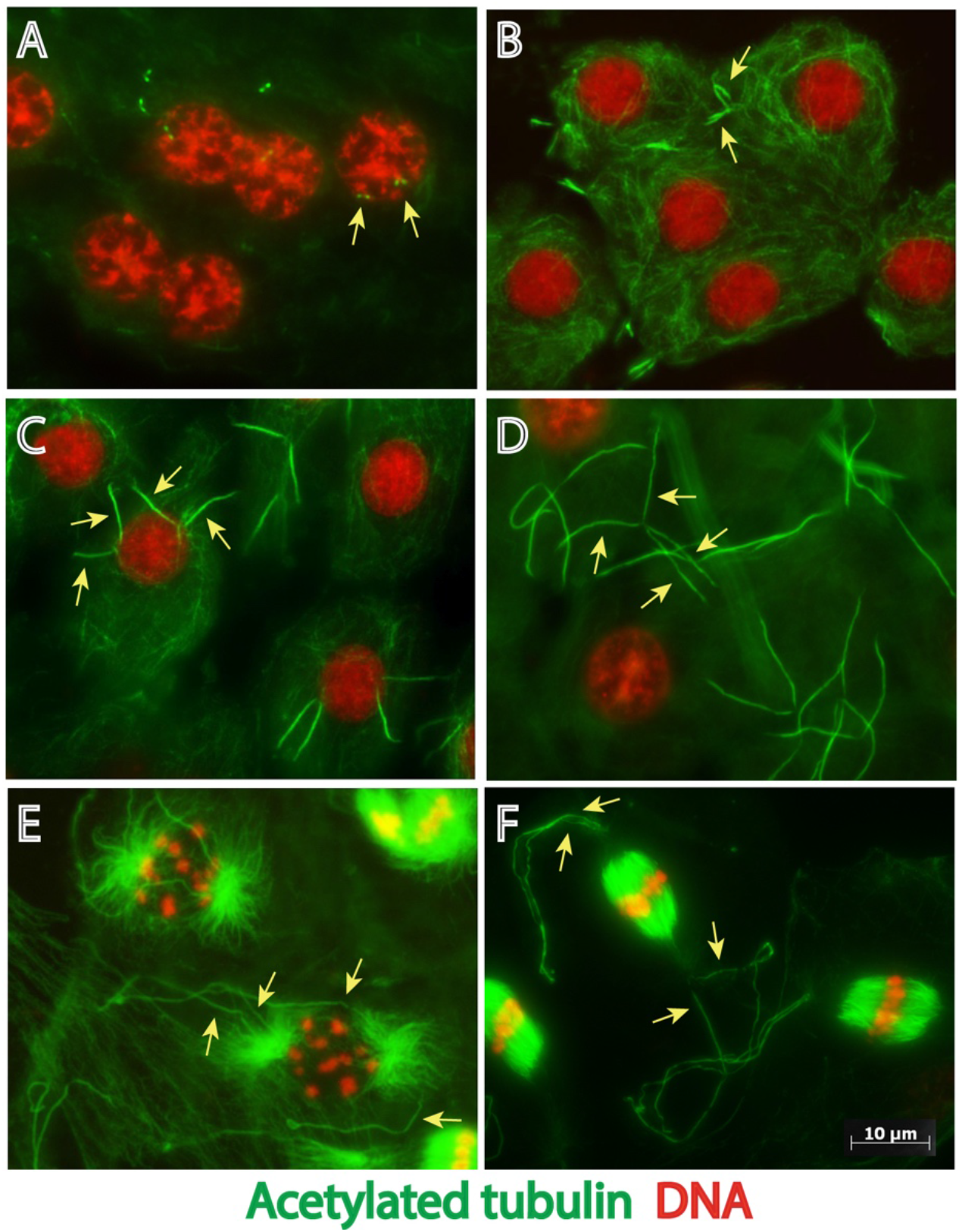
Stages of cilium assembly in *P. brassicae* spermatocytes during meiotic progression. Primary spermatocytes containing two pairs of centrioles in prophase (arrows in A) assemble two pairs of cilia (arrows in B-F) that elongate during progression of the first meiosis. Images of spermatocytes during (A,B) Early prophase, (C) Mid-prophase, (D) Late prophase, (E) Prometaphase, (F) Metaphase. Green: acetylated microtubules, red: DNA. Scale bar: 10 µm

### Distinct ultrastructural features accompany cilium assembly in P. brassicae spermatocytes

To better understand the structural organization of *P. brassicae* spermatocyte cilia we analyzed them by electron microscopy (EM). Ultrastructural sections of early primary prophase spermatocytes showed newly duplicated centriole pairs that reside near the Golgi apparatus (Fig. 2A). During leptotene, the two pairs of orthogonally arranged centrioles had moved to the cell periphery, docked at the plasma membrane, and converted into BBs as it appears that cilium assembly was initiated (Fig. 2B). Docking to the plasma membrane was coincident with the appearance of a thin layer of electron-dense material at the membrane distal to the BB and a small bud formed at the cell surface (Fig. 2B,C). Outgrowth of the cilia began in zygotene with the formation of short ciliary projections containing a growing axoneme (Fig. 2D). In pachytene stage, the cilia, while still short, underwent a significant morphological change and their distal tips swelled (Fig. 2E). Thin filamentous structures (TFS) of electron-dense material were present just below the membrane of the growing cilium and extended outward along the plasma membrane (Fig 2F). The TFS filaments are configured along the length of the short, growing cilium and splay outward from the cilium at its base (Fig. 2F). Grazing sections show that TFS are arranged in parallel rows that run just below the ciliary membrane (Fig. 2G). Higher magnification of the basal region of the cilia revealed regularly spaced links connecting the ciliary membrane and the TFS (Fig. 2H). Remarkably, thin projections link the distal region of the centrioles with the TFS running below the ciliary membrane in newly forming cilia (Fig. 2I). We propose that these TFS structures, which are not seen in Drosophila spermatocytes, are long preformed IFT trains.

**Figure 2.**
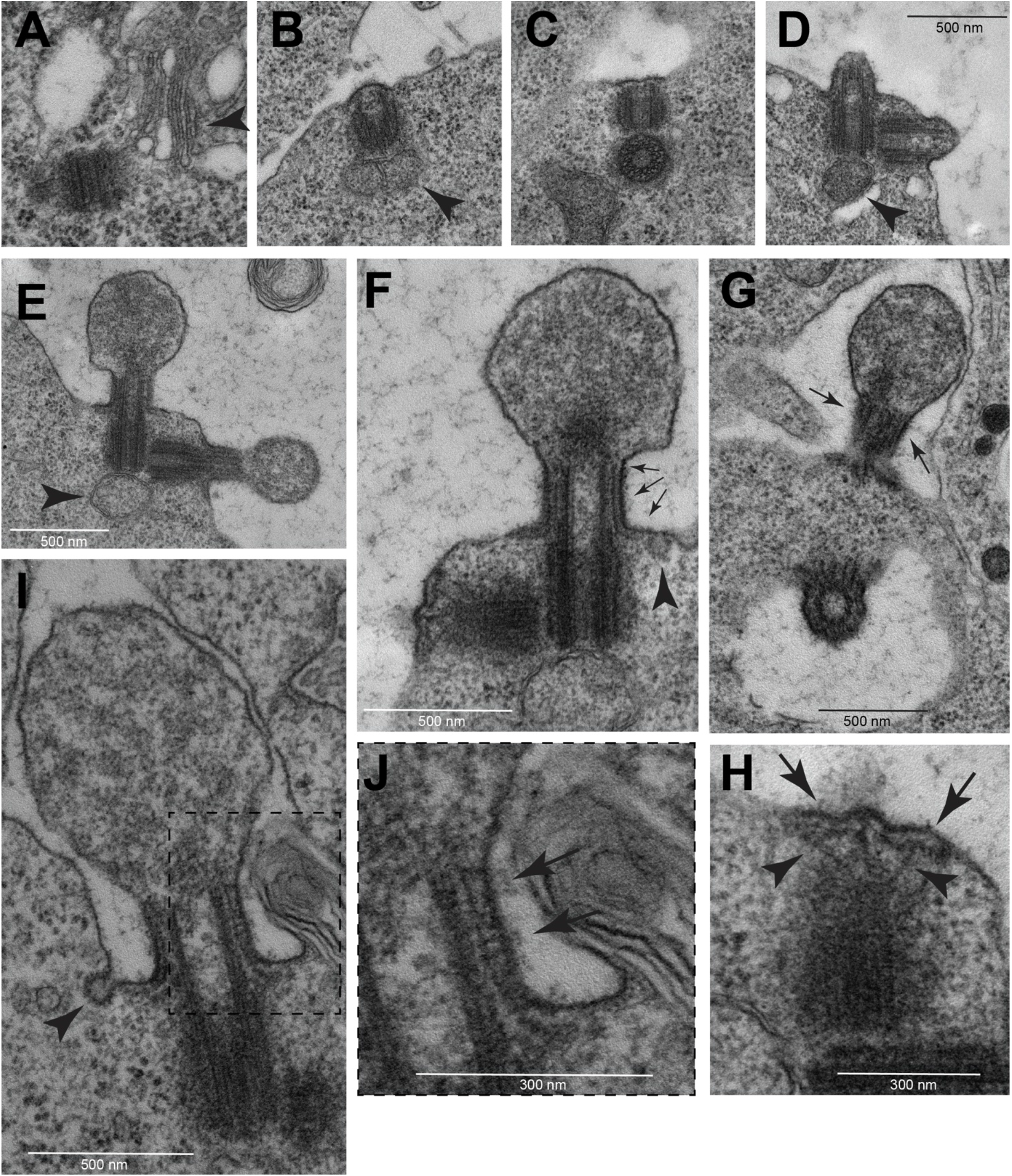
TEM images of early assembly stages of spermatocyte cilia. (A) At beginning of prophase the centrioles are located near the nucleus in association with Golgi cisternae (arrowheads). During leptotene the centrioles dock to the cell membrane (B) and form small buds (C) that elongate progressively during zygotene (D). During pachytene the distal region of the elongating cilia swell (E). One small mitochondrion is associated with the base of the mother centrioles (arrowheads in B,D,E). (F) L-shaped filamentous structures run from the base of the ciliary membrane toward the distal region of the cilium (arrows). (G) Grazing section of the CLR showing parallel bundles of these L-shaped filamentous structures (arrows). (H) Detail of the daughter centriole seen in (F) showing the sub membranous filaments (arrows) and distinct radial filaments emerging from the apical region of the centriole (arrowheads). (I) Early stage cilium and detail of its distal region (boxed region shown in J) showing regularly spaced links (arrows) connecting the ciliary membrane and the filamentous structures. Small vesicles are found at the base of the early cilia close to the cytoplasmic ends of the filamentous structures (arrowheads in F,I).

### Dynamic changes occur at basal bodies and cilia during meiotic prophase

To analyse changes to the structure of BBs, transition zones and cilia at different stages of prophase we prepared serial cross sections from primary spermatocytes. At pachytene the BB consisted of nine MT triplets with a distinct cartwheel and the MT wall was surrounded by an outer layer of thick electron dense material (Fig. 3A). At the distal end of the BB where it transitions to the emerging axoneme of the cilium (the transition zone) the outer layer is no longer visible, and the C-tubules are reduced to hook-shaped sheets (Fig. 3B). Distinct fibres, which appear to correspond to the proximal region of the radial filamentous structures seen in longitudinal sections (Fig 2H), run close to the distal ends of the centrioles (arrows in Fig 3B,F,J).

**Figure 3.**
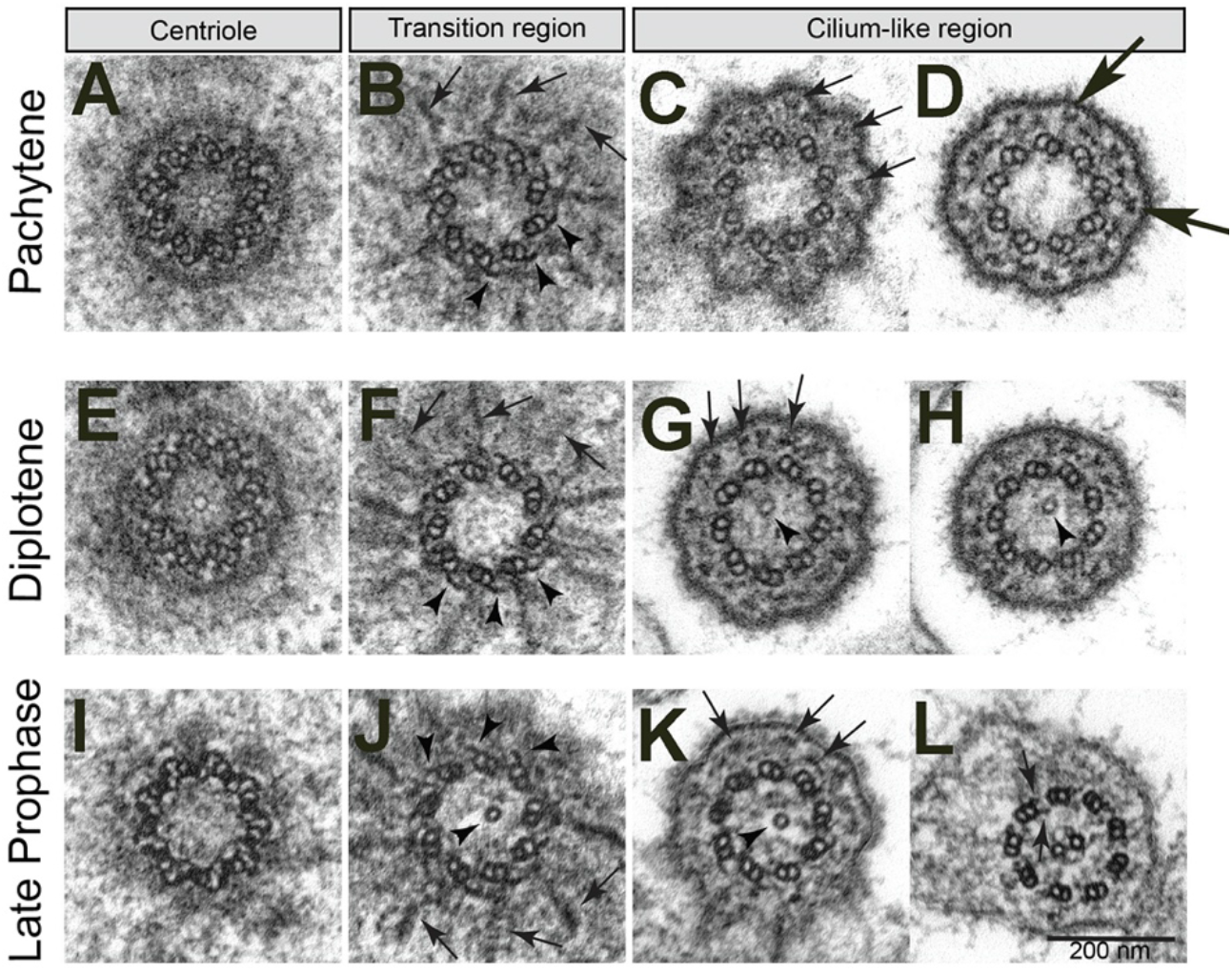
Features of centrioles and cilia during primary spermatocyte meiotic progression. Cross sections during (A-D) pachytene, (E-H) diplotene, (I-L) late prophase. (B,F,J) Radial filamentous structures (arrows) are seen at the transition zone between the distal region of the centrioles and the beginning of the ciliary axoneme; the C-tubules reduced to short hook-like structures (arrowheads). A single tubule is present in the lumen of the basal body and/or transition zone (arrowheads, G,H,J,K). (C,G,K) The filamentous structures running close to the ciliary membrane appear in cross sections as small aggregates of electron-dense material (arrows). (D) Y-linkers connect the A-tubule with the plasma membrane of the basal region of the cilium (large arrows). (L) Late primary spermatocytes display a ciliary axoneme consisting of a canonical 9+2 model with dynein arms (arrows).

The plasma membrane at the base of the cilia bulge out, producing nine small lobes with a flower-like appearance in cross section (Fig. 3C). Small electron-dense aggregates that correspond to cross sections of the filamentous structures running within the basal region of the cilia (Fig. 2F,G) are present in the space between the axonemal MTs, which has a 9 + 0 pattern, and the ciliary membrane (Fig. 3C,D). Strands that appear analogous to transition zone Y-linkers connect the doublet MTs to the ciliary membrane (Fig. 3D). The organization of the BB and the axoneme during pachytene and diplotene did not change (Fig. 3E-F), except that in diplotene a single central MT appeared within the axoneme (Fig. 3G-H). The central tubule extends into the lumen of the BB at the end of the first prophase, when cilia reached their maximum length (Fig. 3I-K). A similar luminal central tubule spanning the basal body and transition zone is present in Drosophila spermatogenesis (Carvalho-Santos et al., 2012; Riparbelli et al., 2013; Riparbelli et al., 2012; Roque et al., 2012; Tates, 1971), and is proposed to promote assembly of the central pair and requires Bld10 for its stability (Carvalho-Santos et al., 2012; Roque et al., 2012). The makeup of the central tubule is not clearly established, but in Drosophila, where the central tubule has been investigated more extensively, taxol treatment resulted in the formation of more than one tubule (Riparbelli et al., 2013), further suggesting that the central tubule is a microtubule. The transition region of the cilium contains nine MT doublets with the remnants of the C-tubules present in the form of hook-like blades. The microtubules of the axoneme have a 9+2 structure that is typical of a motile flagellum. Consistent with a motile architecture, the nine doublet MTs are associated with outer and inner dynein arms (arrows in Fig 3L). A central apparatus, consisting of a central MT pair and nine radial spokes that link the central MT pair to the B-tubule are also consistent with a motile flagellum (Fig. 3L).

### Axoneme elongation

The initial phase of axoneme elongation is evident within the large dilation of the ciliary extension (Fig. 2E). The distal enlarged region of the cilia have enigmatic bulging vesicles with a distinct electron dense membrane (Fig. 4A1,A2). It is unclear if these large vesicles represent fusion or exocytotic events. However, irregular blebs of membrane-bound cytoplasmic material were commonly observed in the intercellular space between primary spermatocytes during cilia growth (Fig. 4B, arrows). Blebs and large cytoplasmic vesicles contained dark small granules that appear to be ribosomes, similar to those seen within the cytoplasm of the spermatocytes (Fig. 4B). These potentially dynamic large vesicles appear to be abundant in the vicinity of spermatocytes (Fig 4B).

**Figure 4.**
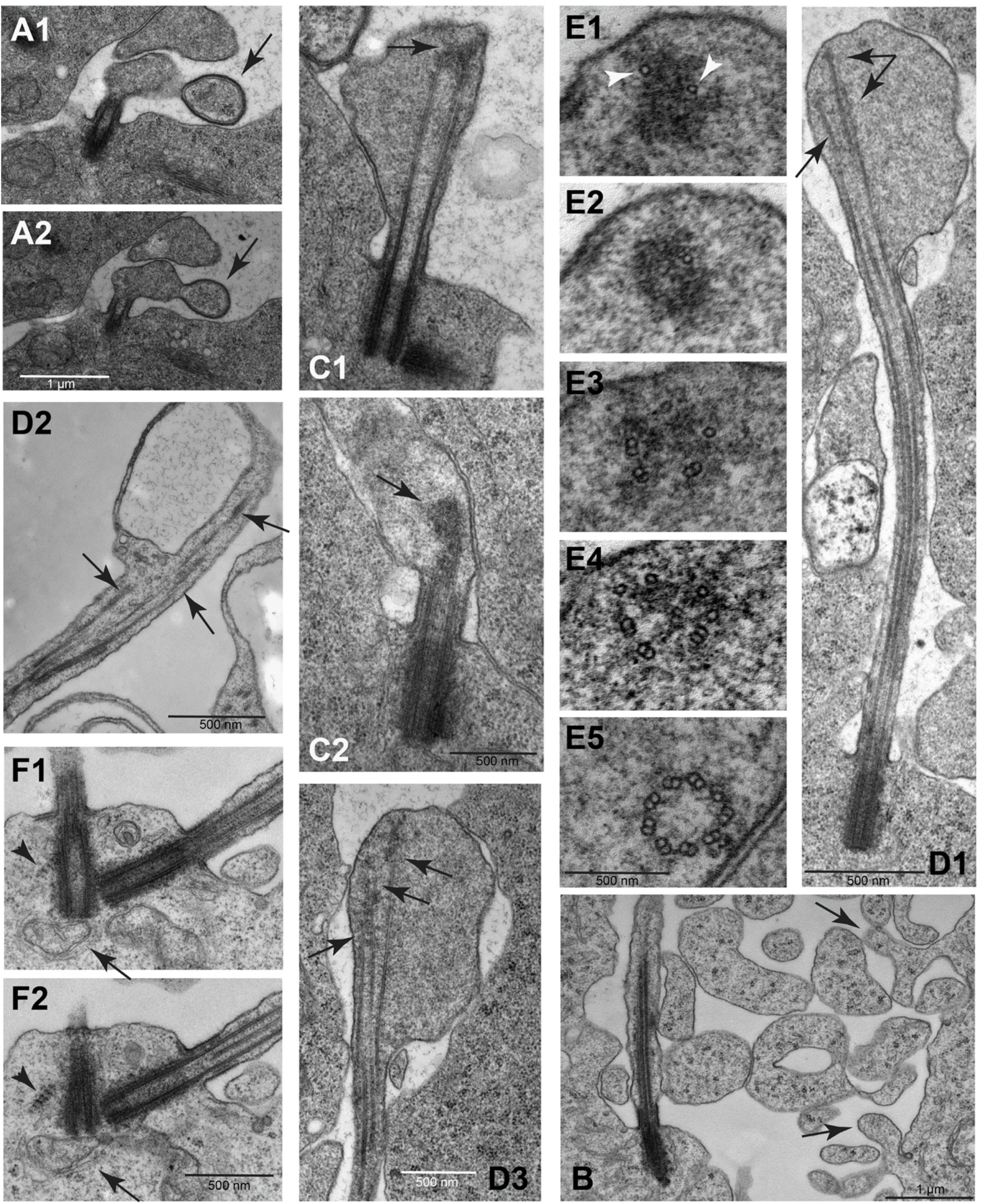
Unique aspects of cilium growth in *Pieris* spermatocytes. (A1-A2): Consecutive sections showing the contact of a large extracellular vesicle (arrow) with the growing cilium. (B): The extracellular space of the spermatocytes during the cilia growth phase is filled with free large vesicles containing ribosomes and distinct cytoplasmic blebs emerge from the cell membrane (arrows). (C1-C2): As prophase progresses the distal ends of the elongating cilia still maintain a distinct swelling where the microtubules end in a distal cluster of electrondense material (arrows). (D1-D3): The microtubules within the distal swellings have variable lengths (arrows); D3 is a close-up of a different section of the sample shown in D1. (E1-E5): Serial sections from the apex of a cilium to its base: single microtubules are associated with the dense material at the distal end of the cilia (arrowheads); moving apically toward the basal body, there are singlet microtubules, then scattered doublets, and at the base of the cilium doublets are arranged in a usual nine-fold symmetry. Central doublets and dynein arms are lacking in early stage cilia. (F1-F2): Striated structures resembling ciliary rootlets are found in close association with the mother centrioles in primary spermatocytes (arrow).

Golgi cisternae were also located close to the centrioles at the base of the cilia (arrow in Fig. 4F). Small vesicles, indicative of active vesicular trafficking at the base of the cilia, are located at the cytoplasmic ends of the TFS filaments (Fig. 2F,I).

Axoneme growth appeared continuous during prophase progression (Fig. 4C,D) while also developing and maintaining a distinct membrane swelling at distal region of the cilia (Fig. 4D). The distal, growing region of the axoneme consists of MTs of different lengths (Fig. 4D1-D3) where the longer MTs end in a cluster of electrondense material (Fig. 4C,D). Serial sections from the apical region of the cilia to its proximal region showed that single MTs were immersed into the dense material at the distal tip (Fig. 4E1,E2). Toward the proximal region single and doublet MTs are formed (Fig. 4E3), then groups of doublet MTs (Fig. 4E4) and finally, near the proximal end, mature doublet MTs arranged in the typical nine-fold symmetry at the axoneme base (Fig. 4E5). Central doublets and dynein arms were lacking in early stage cilia (Fig. 3D,H) but were acquired by the end of prophase (Fig. 3L). Remarkably, the cilia of the mature spermatocytes showed a faint beating movement that we observed during dissection of testis (not shown).

Concurrent with the elongation of the spermatocyte cilia, the centrioles grew in length by about 3-fold, from about 150 nm to 500 nm. In addition, the spermatocyte centrioles grew asymmetrically, with the mother growing longer than the daughter (Fig. 2D,F). Another novel feature of these centrioles are striated structures near the distal end of the basal bodies that appear similar to ciliary rootlets (Fig 4F, arrowheads). However, these structures did not emerge from the proximal base of the centrioles as is typical for ciliary rootlets, but instead appeared to emerge from the distal regions. Moreover, the striated structures were absent during the previous stages of cilium growth.

During the meiotic divisions, the cilia were internalized within deep ciliary pockets and the centrioles organized the centrosomes at the poles of the meiotic spindles (Fig. 5A). Each cilium was then inherited by the spermatids and represented the start point for the growing sperm axoneme that progressively elongated within the cytoplasm (Fig. 5B-D). This maintenance of cilia throughout the division process is exceptional compared to cilium dynamics in typical eukaryotic cells where cilia are resorbed prior to mitosis, but for these spermatocytes is very similar to what happens in Drosophila spermatocytes at meiosis (Riparbelli et al., 2012). Moreover, cilium assembly in spermatids occurs in the cytoplasm, similar to the cytoplasmic growth of axonemes in Drosophila spermatids. At the spermatid stage, the base of the ciliary projection resembles the ‘ring centriole’ of *Drosophila*. However, the ring centrioles of *Pieris* spermatids maintained the filamentous structures (Fig. 5E) that characterized the spermatocyte cilia during the first prophase and the following meiotic divisions, whereas the ring centriole of *Drosophila* displays an accumulation of unstructured electrondense material (Gottardo et al., 2018). In addition, the *Drosophila* axoneme lacks dynein arms within the ciliary cap and dynein arms are assembled onto the axoneme as it assembles in the cytoplasm proximal to the ring centriole. In contrast, cross sections of the ciliary projections of *Pieris* spermatids showed 9+2 axonemes with incorporated dynein arms (Fig. 5D’), as were seen in earlier stages in mature primary spermatocytes (Fig. 3L).

**Figure 5.**
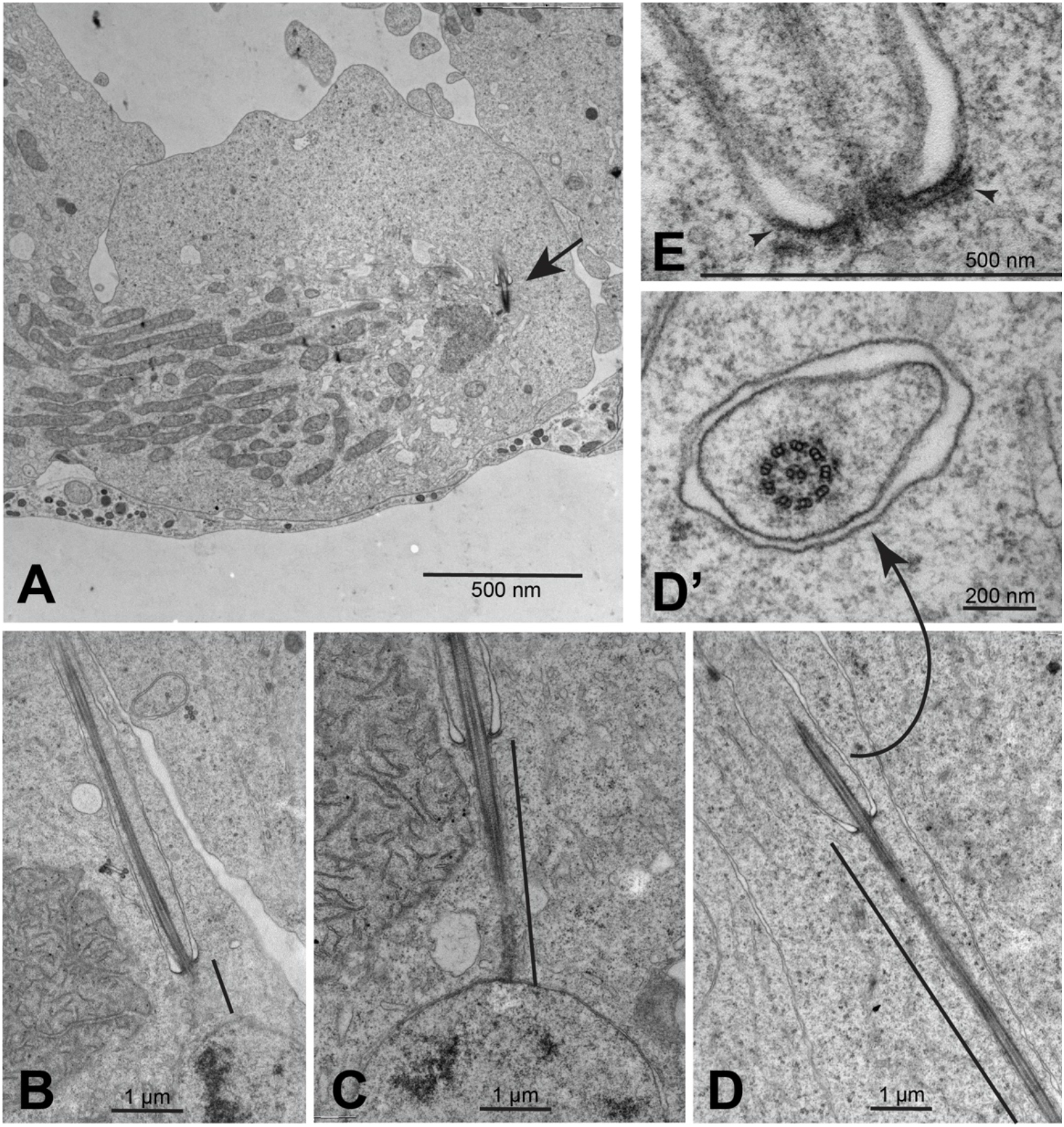
Dynamics of spermatocyte cilia during meiotic divisions. (A) Late anaphase/early telophase of the second meiosis showing the cilium (arrow) in a deep ciliary pocket. (B-D) Early elongating spermatids: the cytoplasmic portion of the axoneme elongates progressively (black line indicates distance of cytoplasmic axoneme elongation). (D’) The axoneme region within the membrane-bound region of the cilium shows a 9+2 structure with dynein arms. (E) The base of the cilia retain the filamentous structures (arrowheads) as those observed in young spermatocytes.

### IFT raft-like arrays in butterfly spermatocyte cilia

Intraflagellar transport (IFT) is an essential process for the growth and maintenance of cilia in most species that was discovered and first described in Chlamydomonas (Pazour and Rosenbaum, 2002; Rosenbaum and Witman, 2002; Sloboda, 2002). IFT complexes are trafficked in clusters or “trains” in the anterograde direction by Kinesin II motors and are remodelled at the cilium tip and then transported in the retrograde direction by Dynein-2. In *Drosophila*, IFT is required for primary cilium assembly, but is dispensable for the assembly of spermatid flagella (Han et al., 2003; Sarpal et al., 2003). Presumably, the cytoplasmic axoneme assembly process does not require IFT, and the short cilium that extends beyond the ring centriole is somehow assembled and maintained independent of IFT. In the butterfly, where the cilia that project beyond the ring centriole are significantly longer than in Drosophila, IFT trains are evident, appearing as electron dense segments adjacent to axonemes in cilia that we interpret as later-stage TFS filaments.

A spacing of about 35 nm separated the ciliary membrane in the basal region of elongating cilia from the axoneme (Fig. 6A). In the distal part of the cilia, this space increased from 45 to 110 nm (Fig. 6B). Longitudinal sections of the proximal region of the elongating cilia showed that apparent IFT trains were present in the narrow space of the ciliary compartment positioned between the axonemal microtubules and the ciliary membrane (Fig. 6A). Therefore, the TFS/IFT trains that appeared in very young primary spermatocytes at the beginning of axoneme formation persisted during prophase progression (Fig 2D). When present at the base of the cilia in young spermatocytes the TFS had similar lengths (Fig 2F). By contrast, the TFS were of variable length at the proximal region of the elongating cilia at the end of the first prophase (Fig 6A).

**Figure 6.**
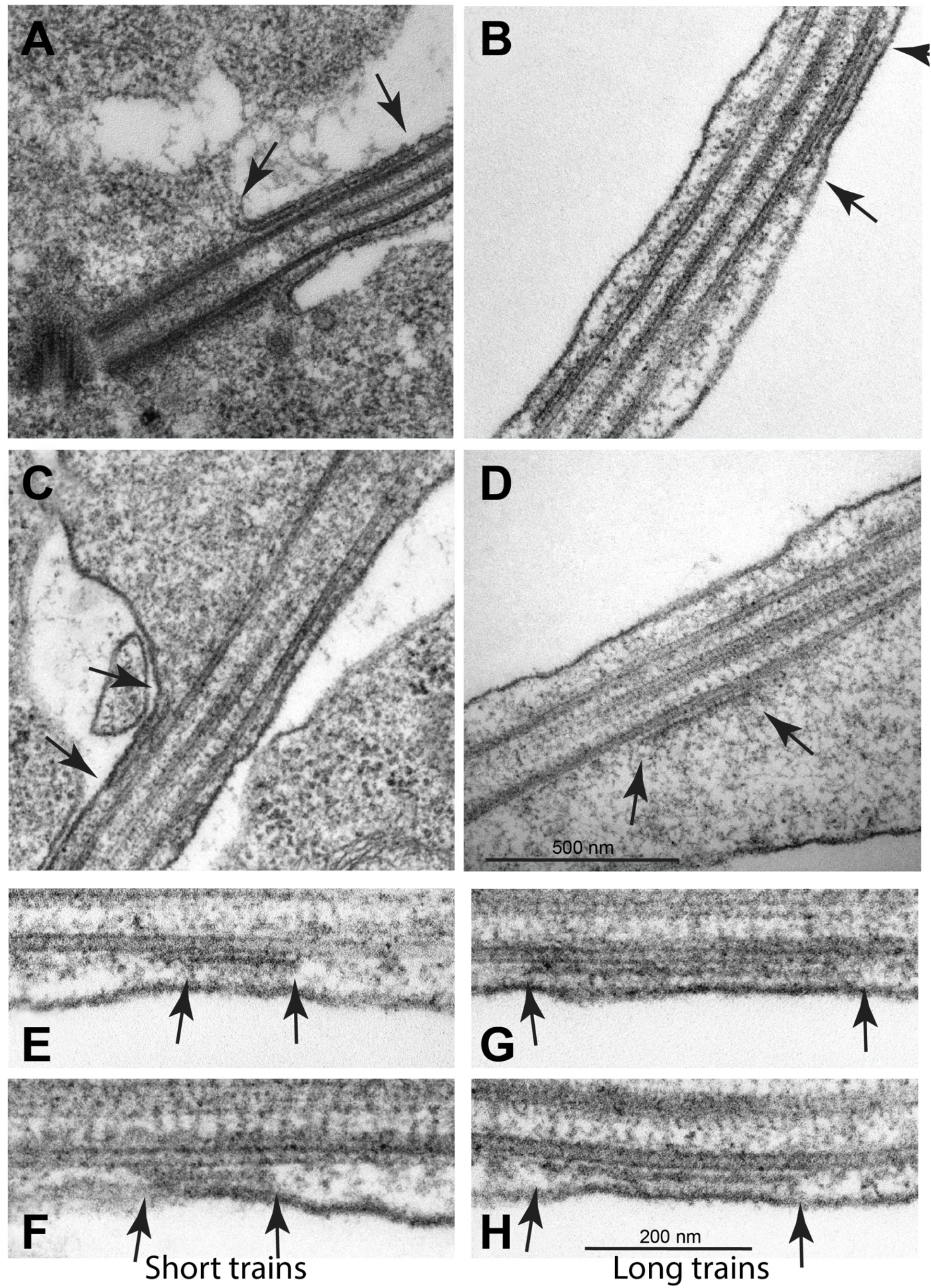
Apparent IFT rafts in *Pieris* spermatocyte cilia. (A) Longitudinal section of the proximal region of a cilium in late primary prophase spermatocytes showing apparent IFT rafts of different lengths (arrows). Filamentous structures that appear to be IFT rafts are present along the cilium (B), at the base of the apical swelling (C) and within the swelling (D) (arrows) always in association with the ciliary axoneme. Longitudinal sections show that there are short (E,F) and long (G,H) raft structures (trains). Thin links connect the raft structures to the ciliary axoneme which might be motor complexes involved in raft transport.

In further support of their interpretation as IFT trains, distinct TFS segments were found associated with the axoneme along the middle of the cilia (Fig. 6B), near the base of the apical swelling (Fig. 6C), and within the apical swelling (Fig. 6D). Within the distal swelling TFS segments retained close contact with the axoneme consistent with their role as IFT trains (Fig. 6D). In longitudinal sections there TFS segments ranged in size from 190-530 nm (Fig. 6E-H). Higher magnification of the TFS revealed faint links connecting both the axonemal MTs and the ciliary membrane (Fig. 6E,F). We propose that TFS are actually IFT trains, and that they form early in spermatocytes.

## Discussion

Insect spermatocytes deviate from the general paradigms of cilium biogenesis and dynamics in several respects. In Drosophila spermatocytes cilia grow from all four centrioles in G2 phase of the cell cycle (Fabian and Brill, 2012; Riparbelli et al., 2012; Tates, 1971), in contrast with the canonical paradigm where cilia form in G1 from only the mother centriole and generally disassemble as the cells enter the cell cycle (Kasahara and Inagaki, 2021; Kim and Tsiokas, 2011; Liang et al., 2016; Plotnikova et al., 2008; Quarmby and Parker, 2005). Here we show that spermatocytes in the butterfly *pieris brassicae* contain motile cilia with evidence of IFT. This contrasts with the best-understood insect system, Drosophila melanogaster, where the spermatocyte cilia are very short and do not require IFT for their assembly nor show evidence of IFT trains even later in spermatid morphogenesis where the flagella grow to a very long length (1800 µm). Moreover, spermatocyte cilia persist throughout the meiotic divisions in Drosophila and in *pieris brassicae* (Gottardo et al., 2014; Riparbelli et al., 2012; Tates, 1971).

IFT is generally necessary for the assembly and maintenance of cilia and flagella (Rosenbaum and Witman, 2002; Scholey, 2003; Sloboda, 2002; Taschner and Lorentzen, 2016). During Drosophila spermatogenesis, however, axoneme assembly occurs within the cytoplasm rather than within a cilium compartment, and so this mode of assembly is presumed to not require IFT and can explain why IFT is not required for flagellum assembly during spermatogenesis. Nevertheless, Drosophila spermatocytes have a short membrane-encased cilium at the distal end of the growing flagellum, but perhaps the short length precludes the need for IFT in its assembly. Therefore it is curious what role the cilia play in spermatocytes, particularly in *pieris brassicae* where the cilia are so prominent and also appear to be motile. One view is that they are assembled in advance of spermatid formation and morphogenesis which begins following two spermatocyte meiotic divisions. However, in Drosophila the cilium partially resorbs in the nascent spermatid, so it does not seem that preassembly accomplishes much if any advantage in this regard. In Drosophila *unc, dila* or *cby* mutants, each of which impairs transition zone assembly, there are no overt defects in meiotic divisions indicating that the cilia have no essential function in the spermatocytes (Baker et al., 2004; Enjolras et al., 2012; Ma and Jarman, 2011). However, in these mutants the morphogenesis of spermatids is impaired and the mature sperm are immotile or mostly immotile despite having apparently normal axonemal ultrastructure in the case of *unc* and *dila* mutants (Baker et al., 2004; Ma and Jarman, 2011). While IFT does not participate in assembly of Drosophila spermatocyte cilia or spermatid flagella, it does appear to function in *pieris brassicae*, and future efforts to investigate the requirement and role of IFT in this and other species might reveal more about the function of these enigmatic cilia in dividing insect spermatocytes.

## Materials and Methods

### Collection and care of Cabbage Butterflies

First instar larvae of the lepidopteran *Pieris brassicae* were collected from cabbage plants in vegetable gardens near Siena Italy and reared in ventilated cages at 20°C. The larvae were feed with cabbage leaves until pupation.

### Immunofluorescence preparations

Testes from fifth instar larvae and pupae were dissected in phosphate buffered saline (PBS). Then the testes were placed in a small drop of 5% glycerol in PBS on glass slides and squashed under a small cover glass and frozen in liquid nitrogen. After removal of the coverslip the samples adhered to the slides were immersed in methanol for 10 min at −20°C. For detection of microtubules the samples were washed for 15 min in PBS and incubated for 1 h in PBS containing 0.1% bovine serum albumin (PBS–BSA) to block non-specific staining and incubated with a mouse anti-acetylated tubulin antibody (Clone 6-11b-1, 1:100; Sigma Cat# T6793) for 4 h at room temperature. After washing for 20 minutes in PBS–BSA the samples were incubated for 1 h at room temperature with a goat anti-mouse secondary antibody conjugated with Alexa Fluor 488 (1:800, Invitrogen). DNA was visualized by incubation of 3–4 min in Hoechst 33342 dye (1µg/ml). Testes were mounted in small drops of 90% glycerol in PBS.

Images were taken using an Axio Imager Z1 (Carl Zeiss, Jena, Germany) microscope equipped with an HBO 50-W mercury lamp for epifluorescence and with an AxioCam HR cooled charge-coupled camera (Carl Zeiss). Gray-scale digital images were collected separately with the AxioVision 2.1 software (Carl Zeiss) and then pseudocolored and merged using Adobe Photoshop 7.0 software (Adobe Systems).

### Transmission electron microscopy

Testes were isolated from pupae and fixed overnight at 4 °C in 2.5% glutaraldehyde in PBS. After rinsing for 30 min in PBS, the samples were post-fixed in 1% osmium tetroxide in PBS for 2 h. After washing for 15 minutes in distilled water the samples were dehydrated in a graded series of ethanol, and then infiltrated with a mixture of Epon–Araldite resin and polymerized at 60 °C for 48 h. Ultrathin sections were cut with a Reichert ultramicrotome, collected with formvar-coated copper grids, and stained with uranyl acetate and lead citrate. TEM preparations were observed with a Tecnai G2 Spirit EM (FEI) equipped with a Morada CCD camera (Olympus).

## Acknowledgements

This research was partially supported by a grant from Ministero dell’Istruzione, dell’Università e della Ricerca to G.C. (MIUR 2020CLZ5XW), and by a National Institutes of Health grant to T.M. (R01 GM139971).

